# Peptide-antibody Fusions Engineered by Phage Display Exhibit Ultrapotent and Broad Neutralization of SARS-CoV-2 Variants

**DOI:** 10.1101/2021.11.29.470362

**Authors:** Jonathan M. Labriola, Shane Miersch, Gang Chen, Chao Chen, Alevtina Pavlenco, Francesca Pisanu, Francesca Caccuri, Alberto Zani, Nitin Sharma, Annie Feng, Daisy W. Leung, Arnaldo Caruso, Gaya K. Amarasinghe, Sachdev S. Sidhu

## Abstract

The COVID-19 pandemic has been exacerbated by the emergence of variants of concern (VoCs). Many VoC mutations are found in the viral spike protein (S-protein), and are thus implicated in host infection and response to therapeutics. Bivalent neutralizing antibodies (nAbs) targeting the S-protein receptor-binding domain (RBD) are promising therapeutics for COVID-19, but are limited due to low potency and vulnerability to RBD mutations found in VoCs. To address these issues, we used naïve phage-displayed peptide libraries to isolate and optimize 16-residue peptides that bind to the RBD or the N-terminal domain (NTD) of the S-protein. We fused these peptides to the N-terminus of a moderate affinity nAb to generate tetravalent peptide-IgG fusions, and showed that both classes of peptides were able to improve affinities for the S-protein trimer by >100-fold (apparent K_D_ < 1 pM). Critically, cell-based infection assays with a panel of six SARS-CoV-2 variants demonstrate that an RBD-binding peptide was able to enhance the neutralization potency of a high-affinity nAb >100-fold. Moreover, this peptide-IgG was able to neutralize variants that were resistant to the same nAb in the bivalent IgG format. To show that this approach is general, we fused the same peptide to a clinically approved nAb drug, and showed that it rescued neutralization against a resistant variant. Taken together, these results establish minimal peptide fusions as a modular means to greatly enhance affinities, potencies, and breadth of coverage of nAbs as therapeutics for SARS-CoV-2.

## Introduction

The COVID-19 pandemic, caused by SARS-CoV-2, has continued to plague the world despite the development of vaccines^1^. To a large extent, this is due to emergent variants of concern (VoCs) that have proven to be more infectious and partially resistant to approved vaccines^2–4^. Consequently, there is an urgent need for alternative therapeutic strategies to complement vaccine campaigns.

SARS-CoV-2 uses its surface spike glycoprotein (S-protein) to interact with host surface receptors and enter host cells. The virus surface displays 25-100 copies of S-protein homotrimers. Each S-protein contains two subunits: the N-terminal subunit (S1) that mediates host cell recognition and the C-terminal subunit (S2) that mediates membrane fusion^5^. The S1 subunit itself contains an N-terminal domain (NTD) followed by a receptor-binding domain (RBD)^6^ that interacts with the host cell-surface protein angiotensin-converting enzyme 2 (ACE2) to initiate infection^7^.

Most natural neutralizing antibodies (nAbs) target the S1 subunit. Many of these bind to the RBD and compete with ACE2^8–14^, but a distinct subset have been shown to target a neutralizing epitope on the NTD^9^. Several natural nAbs have been produced recombinantly and engineered further to develop drugs that have been successful for inhibiting SARS-CoV-2 infection in patients^15^. However, current approved antibody drugs must be administered at very high doses and have proven to be ineffective against many VoCs that have arisen since the original COVID-19 outbreak^2, 4^. Indeed, most VoCs that resist current therapeutic nAbs contain mutations within the RBD that disrupt binding to the nAbs^2^ but not to ACE2^16^. Notably, the B.1.351 variant, which emerged in South Africa and contains critical RBD mutations, is particularly resistant to vaccination and clinical nAbs.

To address these limitations on the potency and breadth of coverage of current bivalent IgG therapies, several groups have developed higher valence protein-based inhibitors^17^. These include small modular Ab domains or non-antibody scaffolds that can be assembled as multimers with enhanced potency due to simultaneous engagement with all three RBDs on an S-protein trimer^18, 19^. Alternatively, we have shown that the fusion of additional Fab arms to either the N- or C-terminus of an IgG heavy chain results in tetravalent IgG-like molecules with enhanced potency and effectiveness against VoCs that resist bivalent IgGs^20^.

Notably, small peptides that target the S-protein with sub-micromolar affinities have been developed and have shown promise as diagnostic tools^21^. Here, we explored whether synthetic peptides that bind to the S-protein could be used to augment the neutralization potency of IgGs in the form of tetravalent peptide-IgG fusions that combine the binding sites of potent nAbs with the small, modular binding sites of peptides. We used naïve phage-displayed peptide libraries to derive synthetic peptides that bind to neutralizing epitopes on the RBD or the NTD. We showed that these small peptides could be fused to a moderate affinity nAb to develop peptide-IgG fusions with affinities enhanced by over two orders of magnitude. Most importantly, one such peptide fusion was able to greatly enhance the neutralization potency against SARS-CoV-2 and VoCs for a high affinity nAb we had engineered earlier^20^, and also, for a clinically approved therapeutic nAb developed by others^10^. Thus, these synthetic peptides hold great promise to enhance the potency and breadth of coverage of therapeutic nAbs against SARS-CoV-2 and its VoCs.

## Results

### Isolation and characterization of S-protein-binding peptides

In order to isolate novel peptides that bind to the S-protein of SARS-CoV-2, we used phage-displayed libraries of 16-residue peptides. We pooled together phage representing a panel of 10 peptide libraries in which diversified positions were encoded by an equimolar mixture of 19 codons representing all genetically-encoded amino acids except cysteine. An “unconstrained” library (X_16_) contained 16 diversified positions with no fixed positions^22^, whereas other libraries contained 14 diversified positions and two fixed cysteine residues separated by 4-12 diversified positions. These “constrained” libraries were designed to display peptides containing disulfide-bonded loops, which have been found to promote tertiary structures that can enhance binding to proteins^26^.

Phage representing the pooled libraries were cycled through five rounds of binding selections with immobilized S-protein ECD or RBD, and several hundred clones were analyzed for binding to the ECD. Clones that exhibited strong binding signals in phage ELISAs with the ECD and negligible signals with bovine serum albumin (BSA) and neutravidin (NAV) were subjected to DNA sequence analysis. This process yielded 160 and 128 unique peptide sequences from the ECD and RBD selections, respectively. Alignment of the sequences revealed that most (85%) of the ECD-selected peptides were derived from two libraries: 63% and 22% were from the X_16_ or C-X_4_-C library, respectively (Figure 1A). In contrast, most (69%) of the RBD-selected peptides were from two different libraries: 42% and 27% were from the C-X_10_-C or C-X_12_-C library, respectively (Figure 1B). Inspection of the aligned sequences revealed significant homology within each peptide family, which enabled us to derive consensus motifs (Figure 1). Notably, the peptides derived from the C-X_10_-C and C-X_12_-C libraries exhibited similar consensus motifs, suggesting that they bind to the S-protein in a similar manner.

**Figure 1.**
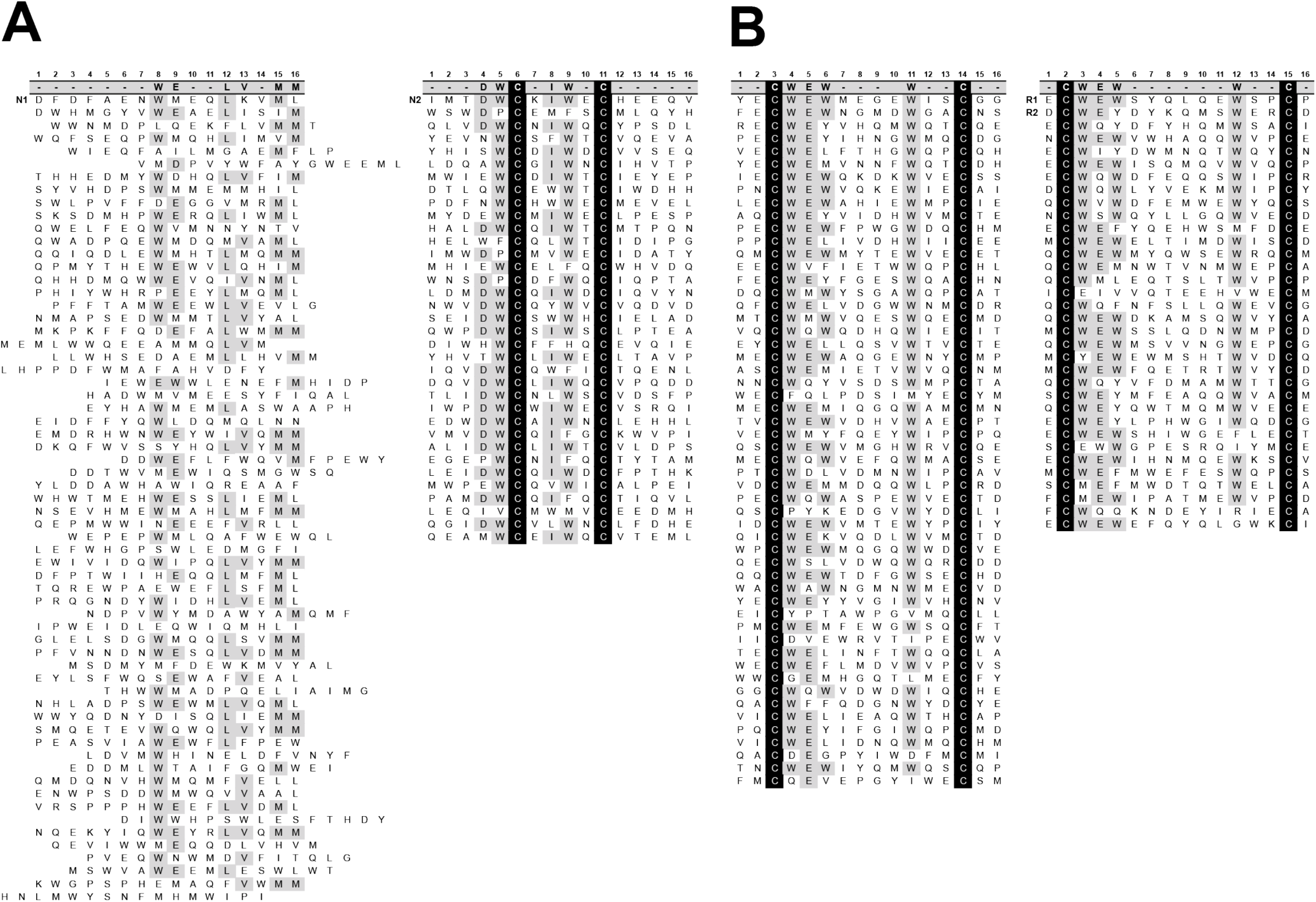
Sequence alignments of phage-derived S-protein-binding peptides. (A) Peptides selected for binding to the S-protein ECD originated from either the X_16_ library (*left*) or the C-X_4_-C library (*right*). (B) Peptides selected for binding to the S-protein RBD originated from either the C-X_10_-C library (*left*) or the C-X_12_-C library (*right*). The positions are numbered at the top, followed by the consensus sequence for positions that exhibited high sequence conservation (>25% for X16, >50% for others), and the unique selected sequences are aligned below the numbering and consensus. Sequences that match the consensus are shaded grey and fixed cysteines are shaded black. Peptides that were characterized in detailed (N1, N2, R1, R2) are labeled and shown at the top each alignment.

For more detailed characterization, we chose four peptides that closely matched the consensus motifs for their respective families. These included a peptide from the X_16_ library (N1) and a peptide from the C-X_4_-C library (N2), which were selected for binding to the ECD, and two peptides (R1 and R2) from the C-X_12_-C library, which were selected for binding to the RBD (Figure 1). As expected, phage ELISAs showed that all four peptide-phage bound to immobilized ECD, but only peptides R1 and R2 bound to immobilized RBD (Figure 2A). To further define where each peptide bound, we assessed whether binding of peptide-phage to the ECD could be blocked by nAbs that recognized known epitopes on either the N-terminal domain (NTD) (IgGs 5-24 and 4-8)^9^ or the RBD (IgGs 15033-7 and REGN10933)^10, 20^. Binding of peptide-phage N1 and N2, but not R1 and R2, was blocked by the two antibodies that bound to a neutralizing epitope on the NTD but not by those that bound to the ACE2-binding site of the RBD. Conversely, binding of peptide-phage R1 and R2, but not N1 and N2, was blocked by the two antibodies that bound to the RBD but not by those that bound to the NTD (Figure 2B). Taken together, these results showed that peptides N1 and N2 likely bind to the NTD whereas peptides R1 and R2 bind to the RBD, and all peptides likely bind to sites that overlap with epitopes of nAbs.

**Figure 2.**
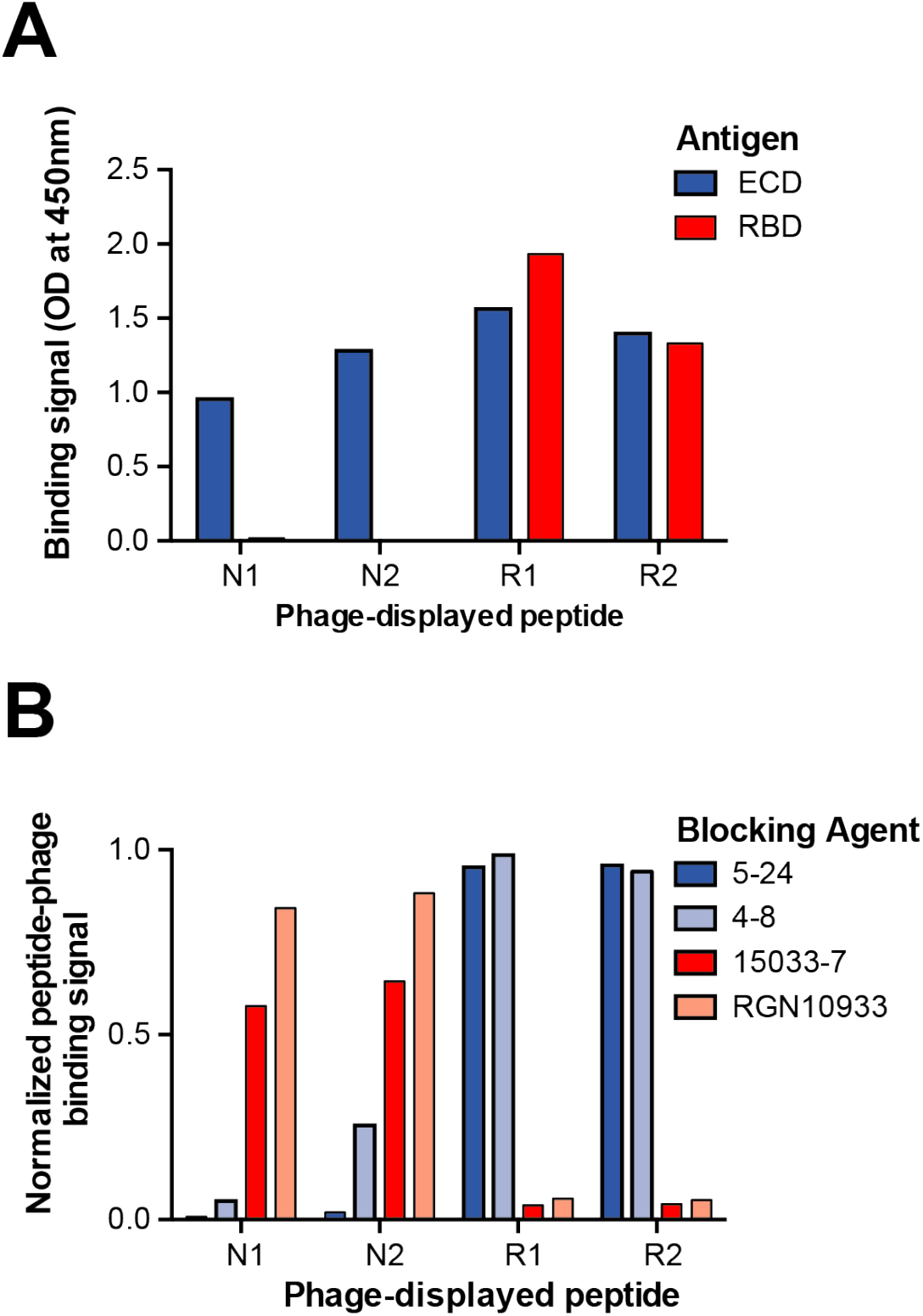
Characterization of epitopes for phage-displayed S-protein-binding peptides. (A) Phage ELISAs for peptide-phage binding to immobilized S-protein ECD (blue bars) or RBD (red bars). Binding was quantified by optical density at 450 nm (y-axis) for 0.5 nM phage displaying the indicated peptide (x-axis). (B) Phage ELISAs for peptide-phage (x-axis) binding to immobilized S-protein ECD in the presence of saturating concentrations of antibodies that bind to the NTD (5-24, dark blue; 4-8, light blue) or RBD (15033-7, dark red; RGN10933, light red). The binding signal (y-axis) was normalized to binding in the absence of antibody.

### Optimization of S-protein-binding peptides

We took advantage of the large families of related sequences to design biased peptide-phage libraries to further optimize peptides N1 and R1 (Figure 3). For each peptide, we used the sequence logo of its family members to identify highly conserved positions that we fixed as the wt, and variable positions for which we used degenerate codons that encoded for the wt sequence and other sequences that were prevalent in the sequence logo. Phage pools representing each biased library were cycled through five rounds of selection for binding to the S-protein ECD, positive clones were identified by phage ELISA, and DNA sequencing revealed 86 and 25 unique sequences from the N1 and R1 libraries, respectively (Figure 4). We used the unique sequences to derive a sequence logo for each selection pool (Figure 3). We chose unique peptide sequences that closely matched the logo for further characterization, reasoning that these likely represented high affinity binders.

**Figure 3.**
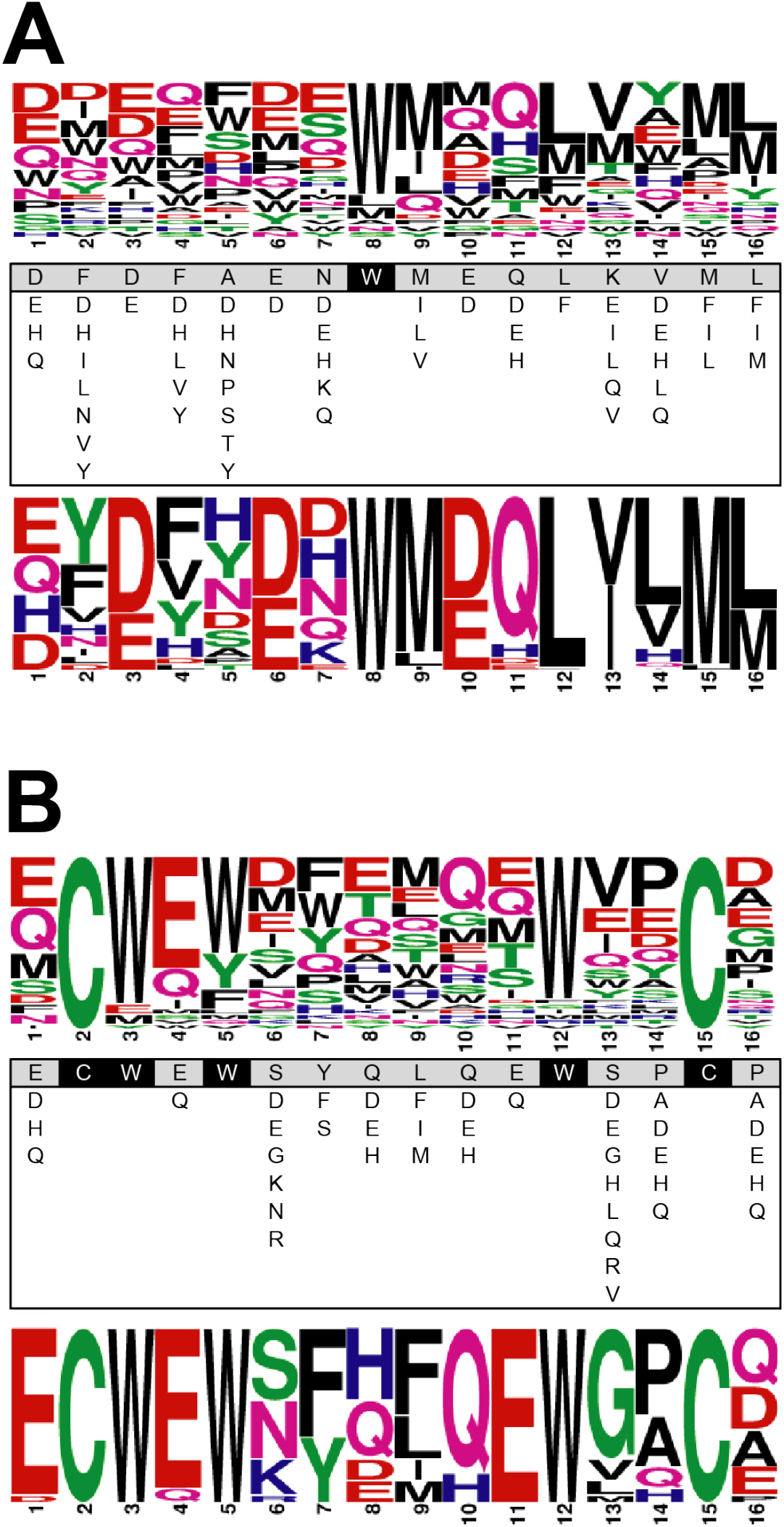
Optimization of S-protein-binding peptides. Biased peptide-phage libraries were designed to optimize (A) peptide N1 or (B) peptide R1. For each peptide, the sequence logo at the top was derived from the family of peptides selected from naïve peptide-phage libraries (Figure 1). Below the logo, the parent peptide sequence is shown with positions that were fixed or diversified shaded black or grey, respectively. For each diversified position, additional allowed sequences are shown below the parent sequence. The sequence logo at the bottom was derived from peptides from the biased library that were selected for binding to the S-protein ECD (Figure 4).

**Figure 4.**
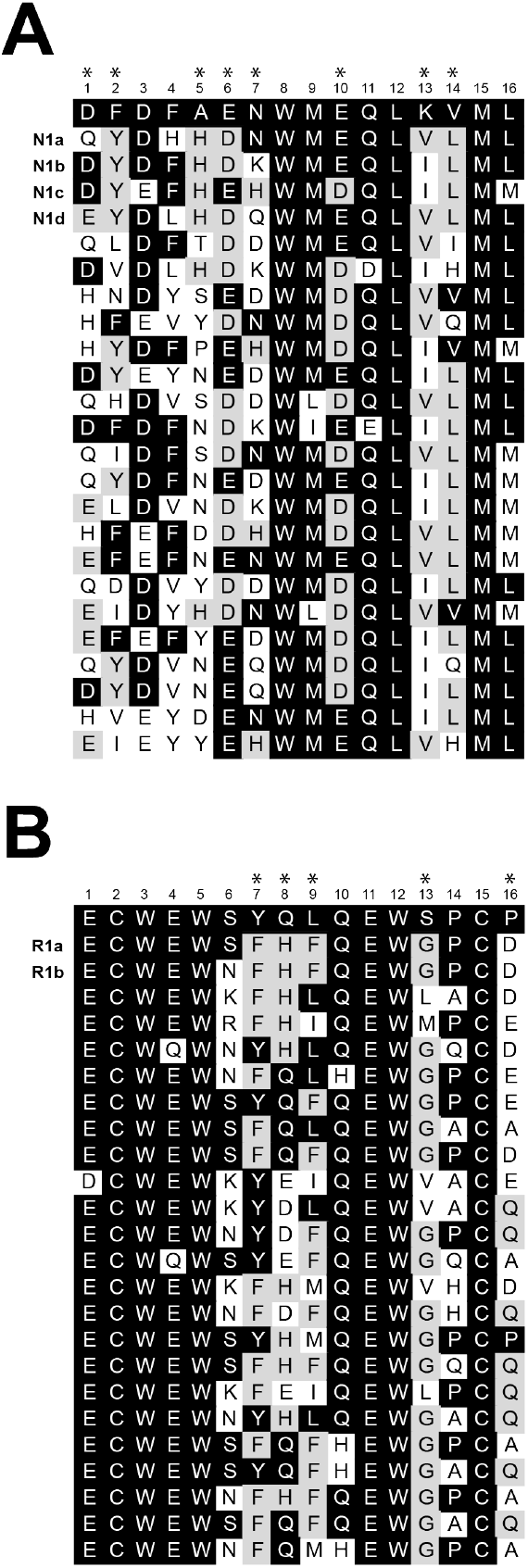
Sequence alignments of S-protein-binding peptides derived from biased phage-displayed libraries. Peptides selected for binding to the S-protein ECD originated from the biased library based on the sequence of (A) peptide N1 or (B) peptide R1 (see Figure 3). Only a subset of peptides from the N1 selection is shown. The positions are numbered at the top, followed by the sequence of the parent peptide. Sequences that match the parental sequence at each position are shaded black. Sequences that do not match the parental sequence but are most prevalent at each position are shaded grey. Asterisks indicate positions at which sequences diverged from the parental sequence, suggesting that sequence differences here may enhance affinity of the variants relative to the parent. The sequences of peptide variants that were chosen for chemical synthesis and further characterization are labeled and shown at the top of each alignment.

Based on this process, we identified four variants of peptide N1 (N1a, b, c, d) and two variants of peptide R1 (R1a, b) for chemical synthesis (Figure 4), and these were synthesized as 22-residue peptides consisting of the following: a Gly residue, followed by the selected 16-residue sequence, followed by a 5-residue hydrophilic linker (Gly-Gly-Lys-Gly-Lys), followed by biotin. We used biolayer interferometry (BLI) assays to determine binding kinetics and affinities. Biotinylated peptides were immobilized on sensor chips that were incubated with solutions of S-protein ECD at various concentrations. Peptide N1 did not exhibit detectable binding, presumably due to low affinity, but its variants bound with moderate apparent affinities (apparent K_D_ = 26-68 nM). Peptide R1 and its variants exhibited high apparent affinities in the single-digit nanomolar range (apparent K_D_ = 0.8-5.6 nM) (Table 1). We note that polyvalent avidity effects likely enhance these apparent affinities, and the intrinsic monovalent affinities are likely lower. Nonetheless, this format does mimic the polyvalent nature of the virus-neutralizing molecules we ultimately intended to produce, as described below.

**Table 1.**
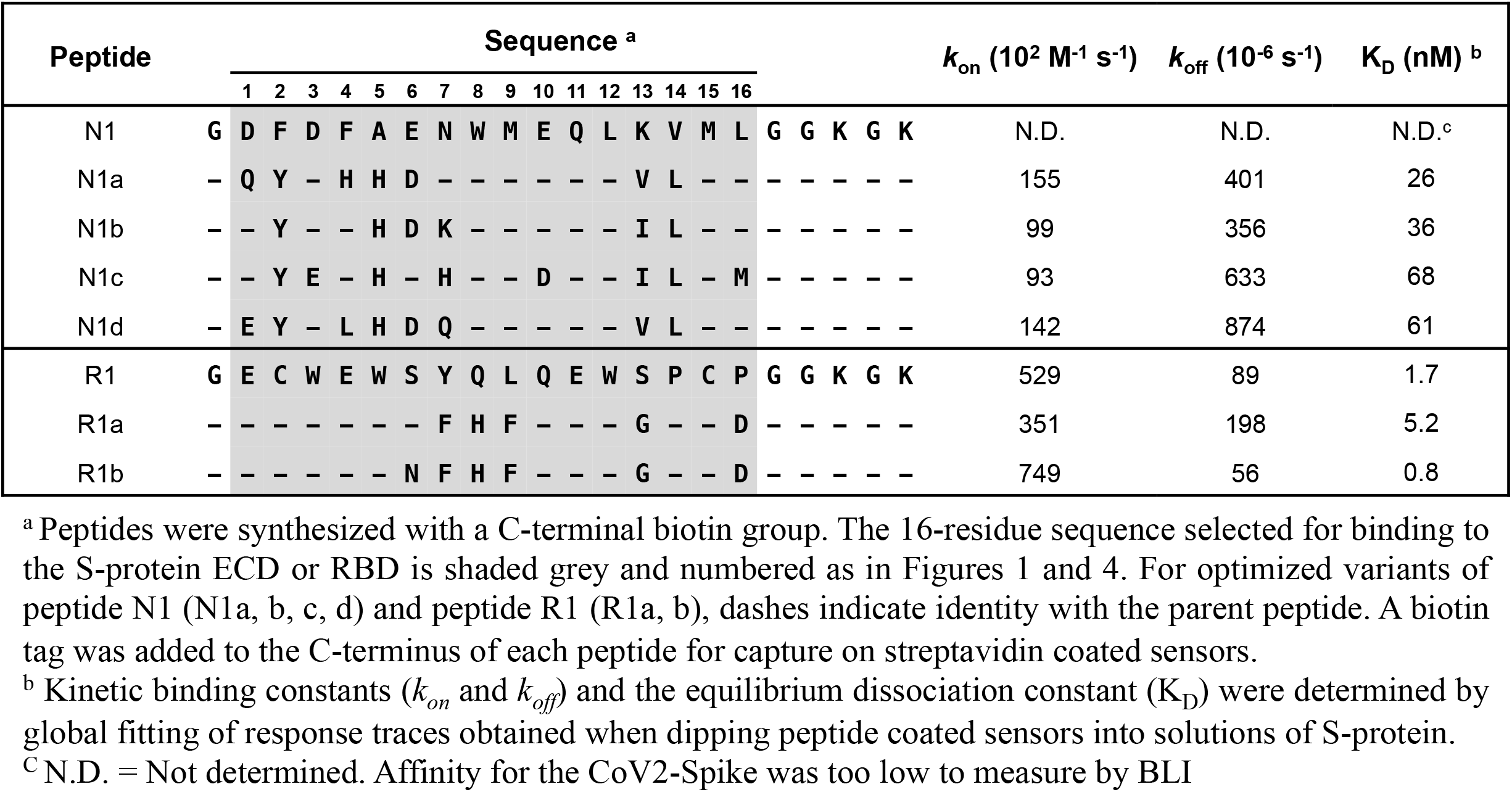
Kinetics of synthetic peptides binding to S-protein ECD.

### Production and characterization of peptide-IgG fusions

We next explored whether peptide fusions could enhance the affinity of a neutralizing IgG targeting the ACE2-binding site of the RBD. For this purpose, we used the moderate affinity IgG 15033 that we had selected from a naïve phage-displayed synthetic antibody library^20^, in order to accurately discern affinity differences. Peptide N1 or R1 was fused to the N-terminus of either the light chain (LC) or heavy chain (HC) of IgG 15033 with an intervening 20-residue Gly/Ser linker. The resulting peptide-IgG fusion proteins and IgG 15033 were purified by transient transfection in mammalian Expi293F cells^20^. All proteins could be purified to near homogeneity by affinity chromatography with protein-A resin, as evidenced by SDS-PAGE (Figure 5A). As expected, under non-reducing conditions, the single bands for the intact peptide-IgG molecules migrated slightly slower than the band for IgG 15033. Under reducing conditions, either the HC band or the LC band migrated slower for peptide-IgG fusions compared with IgG 15033, as expected for HC or LC peptide fusions, respectively. Like IgG 15033 and the highly specific IgG trastuzumab, the four peptide-IgG fusions did not bind to seven immobilized, heterologous proteins that are known to exhibit high non-specific binding to some IgGs (Figure 4B), and lack of binding in this assay is a predictor of good pharmacokinetics *in vivo*^27^.

**Figure 5.**
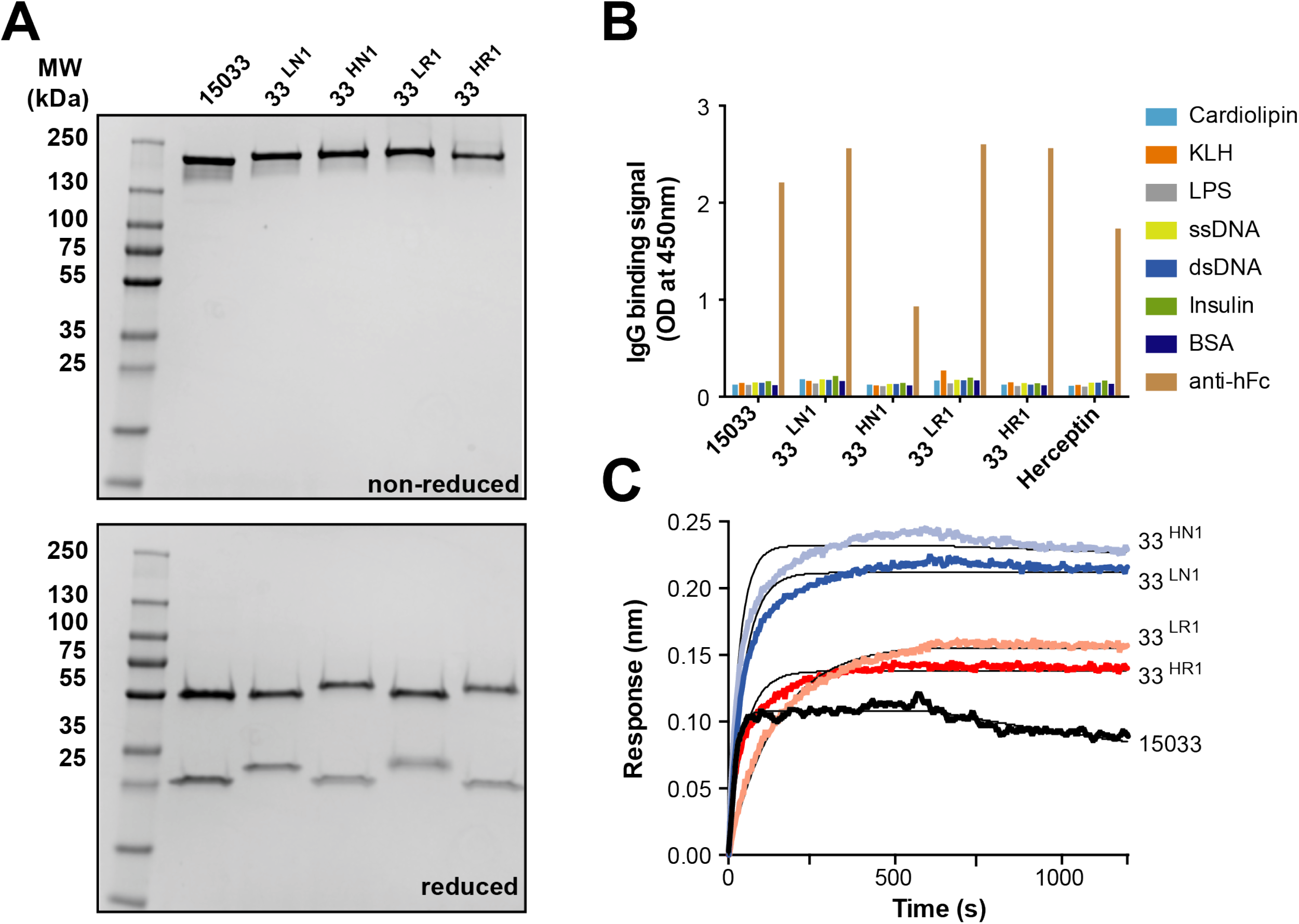
Characterization of peptide-IgG fusion proteins. Peptides were fused to the N-terminus of the HC or LC of IgG 15033, and the resulting peptide-IgG fusions were named as follows: N1 fused to HC, 33HN1; N1 fused to LC, 33LN1; R1 fused to HC, 33HR1; R1 fused to LC, 33LR1. (A) SDS-PAGE analysis of peptide-IgG fusion proteins under non-reducing (top) or reducing conditions (bottom). (B) Assessment of non-specific binding of peptide-IgG fusion proteins to immobilized antigens or a goat anti-human Fc Ab (positive control). (C) BLI sensor traces for 20 nM IgG 15033 or peptide-IgG fusions binding to immobilized S-protein ECD. Association and dissociation kinetics were monitored for 600 seconds each. The derived binding constants are shown in Table 2.

**Table 2.** Kinetics of peptide-IgG fusion proteins binding to S-protein ECD.

| **IgG <sup>a</sup>** | **$k_{on}$ ($10^4 \text{ M}^{-1} \text{ s}^{-1}$)** | **$k_{off}$ ($10^{-6} \text{ s}^{-1}$)** | **$K_D$ (pM) <sup>b</sup>** |
| --- | --- | --- | --- |
| 15033 | 170 | 500 | 300 |
| 33<sup>HN1</sup> | 160 | 38 | 24 |
| 33<sup>LN1</sup> | 110 | <0.1 | <1 |
| 33<sup>HR1</sup> | 120 | <0.1 | <1 |
| 33<sup>LR1</sup> | 35 | 2.2 | 6.1 |

<sup>a</sup> Peptide-IgG fusion proteins consisted of phage-derived peptides fused to the N-terminus of the light chain or heavy chain of IgG 15033. The following nomenclature was used: L = light chain fusion, H = Heavy chain fusion, N1 = NTD-binding peptide 1, R1 = RBD-binding peptide 1 (see Table 1 for sequences).
<sup>b</sup> Kinetic binding constants ( $k_{on}$ and $k_{off}$ ) and the equilibrium dissociation constant ( $K_D$ ) were determined by global fitting of response traces obtained when dipping S-protein coated sensors into solutions of indicated IgG.

Kinetic binding analysis by BLI showed that the tetravalent peptide-IgG fusions exhibited greatly reduced off-rates compared with IgG 15033, and consequently, affinities were greatly improved (Figure 4C, Table 2). These results were as expected, since the addition of a peptide ligand to an RBD-binding IgG should increase the overall enthalpic energy of binding, which in turn should reduce the off-rate. In particular, the peptide-IgG with peptide R1 fused to its HC (33^HR1^) and the peptide-IgG with peptide N1 fused to its LC (33^LN1^) exhibited extremely slow off-rates that were beyond the sensitivity of the instrument, and consequently, the apparent dissociation constants were estimated to be sub-picomolar (apparent K_D_ < 1 pM), which was >100-fold improved relative to IgG 15033 (apparent K_D_ = 100 pM).

### Inhibition of SARS-CoV-2 infection in cell-based assays

Finally, we explored the critical question of whether peptide fusions could enhance the potency of nAbs in cell-based assays of SARS-CoV-2 infection. For this purpose, we fused peptide R1 to the N-terminus of the HC of IgG 15033-7, a more potent variant of IgG 15033 with an optimized LC^20^. The resulting peptide-IgG fusion protein (33-7^HR1^) was compared to IgG 15033-7 in assays that measured the infection of ACE2-expressing Vero E6 cells with a panel of six SARS-CoV-2 variants including the original Wuhan virus (B.1) and later emerging variants of concern (VoCs), including variants isolated in Italy (B.1.1), the United Kingdom (B.1.1.7), South Africa (B.1.351), Nigeria (B.1.525), and Brazil (P.1). In every case, the potency of the peptide-IgG fusion greatly exceeded that of the IgG (Figure 6A). Indeed, peptide-IgG 33-7^HR1^ neutralized three VoCs in the single-digit ng/mL range (IC_50_ = 6.5-7.6 ng/mL) and it neutralized the other three viruses in the double-digit ng/mL range (IC_50_ = 12-49 ng/mL). In contrast, IgG 15033-7 was ineffective against three VoCs (IC_50_ > 900 ng/mL) and was much less effective than IgG 33-7^HR1^ against the other three VoCs (IC_50_ = 64-310 ng/mL).

**Figure 6.**
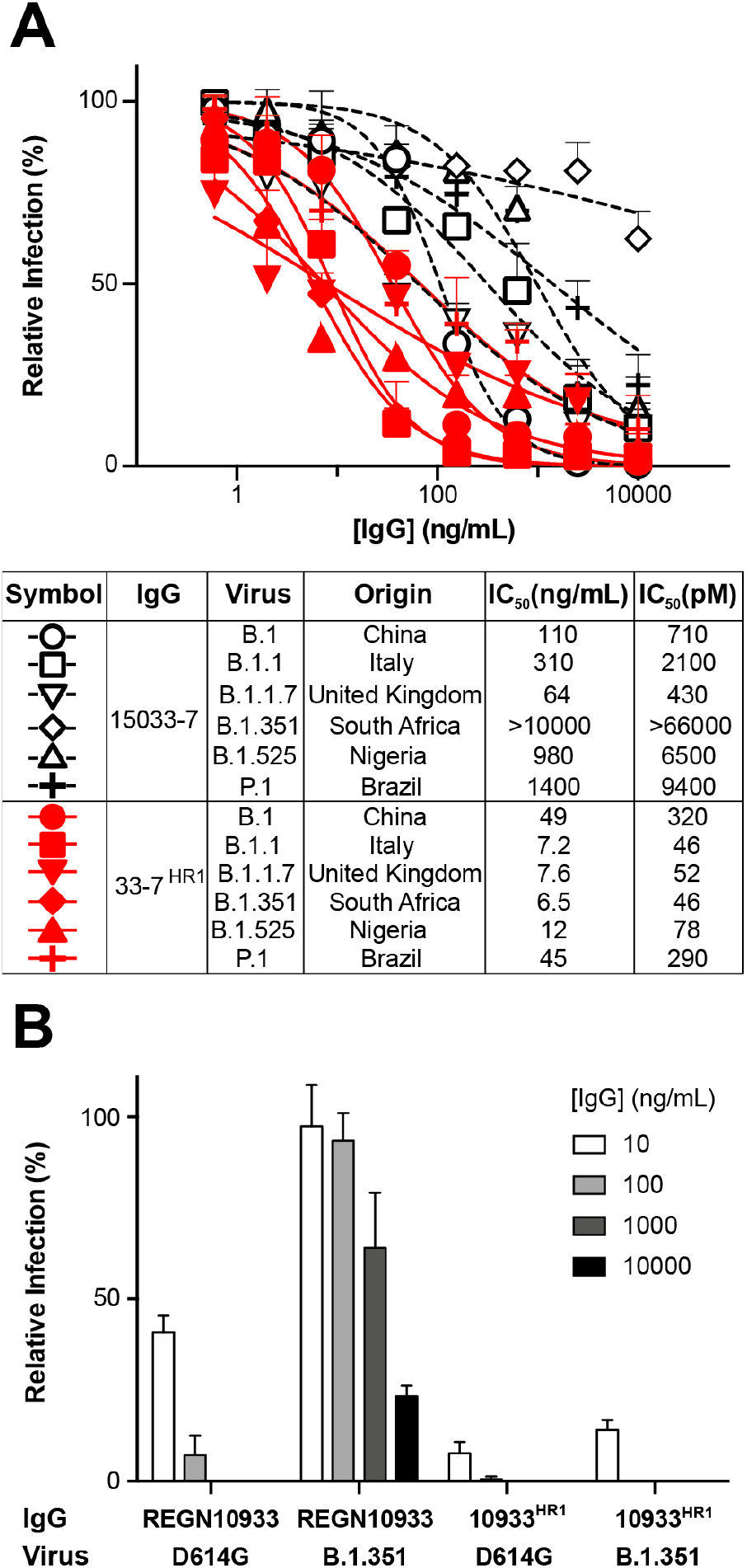
Neutralization of SARS-CoV-2 variants. (A) Authentic virus neutralization assays to test efficacy of IgG 15033-7 and peptide-IgG 33-7HR1 (IgG 15033-7 with peptide R1 fused to the N-terminus of its HC) against isolated SARS-CoV-2 variants. The virus was pretreated with serial dilutions of IgG followed by infection of ACE2-expressing Vero E6 cells measured relative to untreated control. At the top, the graph shows the neutralization curves for six SARS-CoV-2 variants treated with IgG 15033-7 (open black symbols and dashed lines) or peptide-IgG 33-7HR1 (closed red symbols and solid lines). At the bottom, the table shows the symbols used for the curves and the country of origin for each virus variant. IC_50_ values calculated from the curves are shown in ng/mL or nM units. (B) Focus reduction neutralization assays to test efficacy of IgG RGN10933 and peptide-IgG 10933HR1 (RGN10933 with peptide R1 fused to the N-terminus of its HC) against isogenic SARS-CoV-2 variants D641G and B.1.351. The virus was pretreated with serial dilutions of IgG followed by infection of ACE2-expressing Vero E6 cells, which was measured relative to untreated virus. Each value represents the mean of duplicate measurements.

We also explored whether the modularity of the peptides could be exploited to enhance the potency of a clinically approved therapeutic nAb (REGN10933, Figure 6B). We constructed a variant of REGN10933 by fusing peptide R1 to the N-terminus of the HC. The resulting peptide-IgG fusion (10933^HR1^) proved to be more potent than REGN10933 in neutralization assays against the SARS-CoV-2 variant D614G (IC_50_ < 10 or ~10 ng/mL, respectively). Most strikingly, peptide-IgG 10933^HR1^ was also extremely potent against the South African variant B.1.351 (IC_50_ < 10 ng/mL), against which REGN10933 was completely ineffective (IC_50_ >1000 ng/mL), consistent with previous reports^2^. Taken together, these results showed that peptide fusions greatly enhanced the potency and breadth of coverage of neutralizing IgGs against SARS-CoV-2 and its VoCs.

## Discussion

The trimeric structure of the SARS-CoV-2 S-protein can be exploited to engineer Ab-based inhibitors with enhanced neutralization, by engagement of three neutralizing epitopes on a single trimer. Structural studies from us and others have shown that potent neutralizing IgGs engage two RBDs on an S-protein trimer28, and we have shown that the addition of Fab arms to either end of an IgG can further enhance neutralization by enabling engagement to the third RBD20. Here, rather than using large Fabs as the additional binding domain, we used small peptides to greatly enhance the affinities and potencies of tetravalent peptide-IgG fusions compared with our bivalent IgG 15033-7. Most importantly, peptide-IgG 33-7^HR1^ acted as a potent inhibitor of virus variants that resisted IgG 15033-7. Indeed, we showed further that a peptide-IgG version of the approved drug RGN10933 was able to potently neutralize a VoC against which REGN10933 was completely ineffective.

To gain insight into the structural basis for how a peptide within a peptide-IgG fusion could enhance affinity, we examined our published model of two 15033-7 Fabs bound to the S-protein trimer, reasoning that this likely provides an accurate view of how the two Fab arms of a bivalent IgG would bind (Figure 7A). In this model, the RBDs bound to Fabs are in an “up” conformation, whereas the unbound RBD is in a “down” conformation. The C-termini of the two Fab HCs are separated by 45 Å and are well-positioned to be linked to an Fc in an IgG molecule, whereas their N-termini are fairly close to the unbound RBD. In particular, the N-terminus of one HC is 45 Å from the center of the exposed region of the neutralizing epitope for Fab 15033-7 on the unbound RBD. Peptide R1 likely binds to a site that overlaps with this epitope, given that peptide R1 and IgG 15033-7 compete for binding to the S-protein (Figure 2B). Consequently, in peptide-IgG 33-7^HR1^ with two Fabs bound to an S-protein timer, it is reasonable to assume that one of the R1 peptides could bind simultaneously to this third epitope, and the estimated length (~70 Å) of the extended 20-residue linker that connects the peptide to the IgG is consistent with this model.

**Figure 7.**
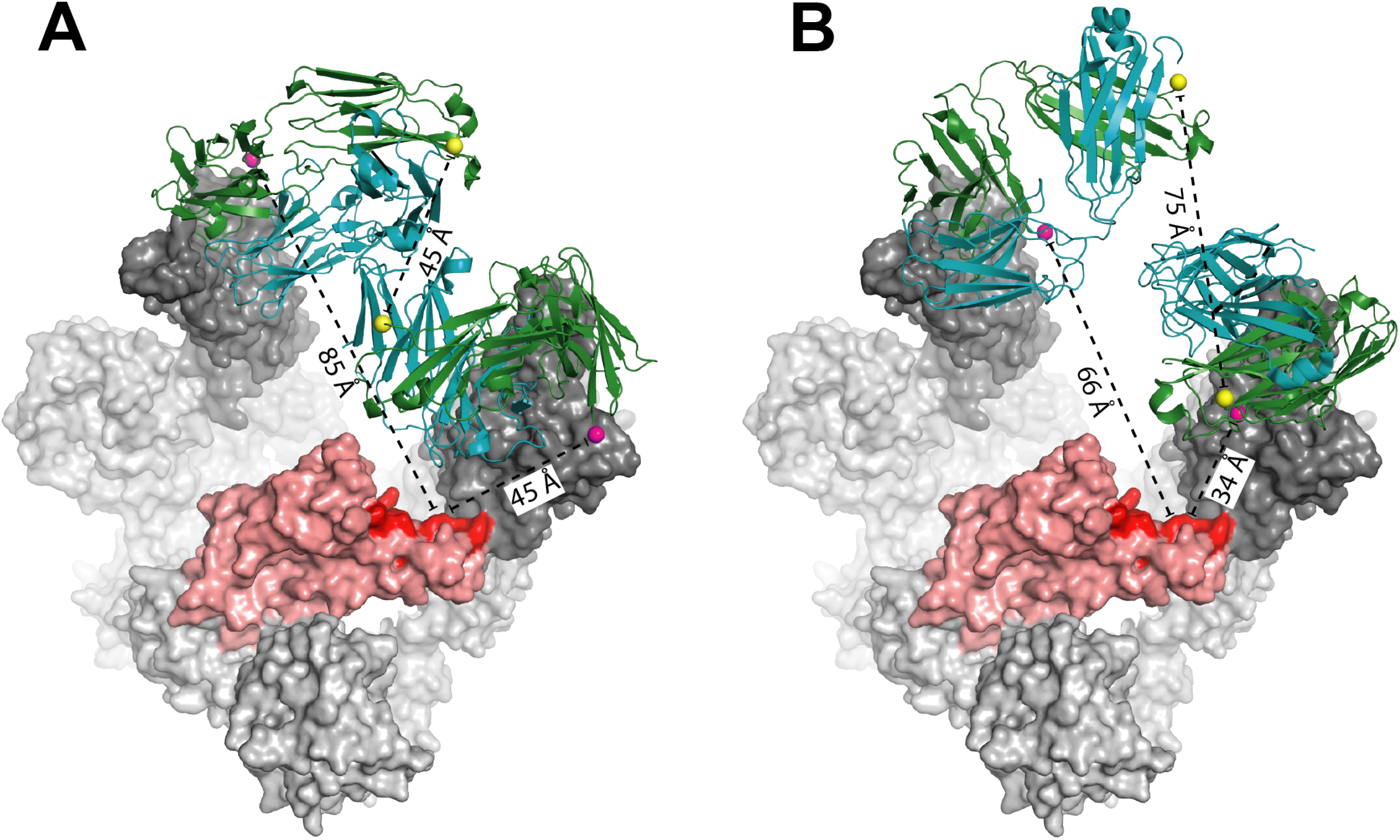
Structural models of multivalent ligands binding to the S-protein trimer. Structural models are shown for (A) two 15033-7 Fabs or (B) two RGN10933 Fabs bound to the S1 subunit of the S-protein of SARS-CoV-2. The S1-protein is shown as a surface colored in light grey, except for the following regions. The bound RBDs in the “up” position are colored dark grey. The unbound RBD in the “down” position is colored dark or light red for residues that are within or outside the epitope for Fab 15033-7, respectively. The Fabs are shown as ribbons with the HC and LC colored forest green or teal, respectively. The C-termini of the Fab HCs are shown as yellow spheres. The C-termini and N-termini of the Fab HCs are shown as yellow or magenta spheres, respectively. Distances are shown as dashed black lines and are labeled accordingly.

We next examined an analogous structural model of two RGN10933 Fabs bound to an S-protein trimer (Figure 7B). The epitopes for REGN10933 and 15033-7 Fabs share significant overlap, but the REGN10933 Fabs are oriented such that the N-termini of the two HCs are 34 Å or 66 Å from the 15033-7 epitope on the unbound RBD. Consequently, the ~70 Å linker that connects each peptide to the IgG would be sufficient to enable either of the two fused peptides within peptide-IgG10933^HR1^ to bind a site close to this epitope.

We also developed peptides that bound to the NTD, likely at a site that overlaps with epitopes for other neutralizing nAbs (Figure 2B), and we showed that peptide N1 greatly enhanced the affinity of peptide-IgG 33^LN1^. While not yet explored, it is likely that the enhanced affinity of peptide-IgG 33^LN1^ should also be reflected in enhanced potency of virus neutralization. Notably, peptides R1 and N1 appear to work best as fusions to the HC or LC, respectively, and given that they recognize completely distinct epitopes, it will be interesting to explore whether dual fusions of the two peptides could enhance the potencies of peptide-IgG fusions even further, and these studies are ongoing.

The ultimate value of peptide-IgG fusions targeting SARS-CoV-2 and its variants resides in the potential for the development of superior therapeutics for the treatment of COVID-19 and other viral diseases. We are currently assessing and optimizing the biophysical properties of the peptide-IgG fusions with the aim of developing drug-grade biologics. However, there is already strong precedent for peptide-IgG fusions as clinical drugs in the form of biologics that have reached phase 2 clinical trials as cancer therapeutics^29^. These anti-cancer biologics are very similar in format to the anti-viral peptide-IgG fusions we report here^30^. Critically, anti-cancer peptide-IgG fusions have exhibited good pharmacokinetic profiles^31^ and tissue penetration^32^, and they have been well tolerated in the clinic^33^. Taken together, our results establish peptide-IgG fusions as a powerful means for greatly enhancing the potency and coverage of next-generation biologics for the treatment of disease caused by SARS-CoV-2 and its variants.

## Materials and Methods

### Protein production

The SARS-CoV-2 S-protein extracellular domain (ECD) and RBD were produced and purified as described^20^ and were a kind gift from Dr. James Rini. Purified proteins were site-specifically biotinylated in a reaction with 200 μM biotin, 500 μM ATP, 500 μM MgCl_2_, 30 μg/mL BirA, 0.1% (v/v) protease inhibitor cocktail and not more than 100 μM of the protein-AviTag substrate. The reactions were incubated at 30 °C for 2 hours and biotinylated proteins were purified by size-exclusion chromatography.

### Antibody production

IgG and peptide-IgG fusion proteins were produced as described^20^. S-protein-binding peptides were fused to the N-terminus of the heavy or light chain through a 20-residue linker (sequence: GGGGSGGGGSGGGGSGGGGS) using standard molecular biology techniques.

### Phage display selections

Naïve libraries were constructed as described^22^. For libraries for peptide optimization, oligonucleotides were synthesized using degenerate codons encoding for the amino acids at each position indicated in Figure 3. Oligonucleotide-directed mutagenesis was performed to introduce randomized sequences fused to the gene-8 major coat protein of M-13 bacteriophage as described^23^. Each phage-displayed peptide library was selected for binding to immobilized S-protein ECD or RBD, as described^23^, but with the following modifications. Biotinylated S-protein ECD or RBD was captured on wells coated with neutravidin (NAV), followed by incubation with the phage-displayed peptide library. After five rounds of binding selections, individual clones were picked and DNA was sequenced. Clones that showed a significant binding signal to S-protein ECD and/or RBD, but not to BSA or NAV, were selected for further analysis.

### ELISAs

Phage ELISAs were performed, as described^20^, but with the following modifications. 384-well maxisorp plates (Sigma) were coated with NAV, or left uncoated as negative control, and blocked with PBS, 0.5% Bovine serum albumin (BSA). Biotinylated target protein was captured by incubation in NAV-coated and BSA-blocked wells, or with buffer solution alone as negative control, at room temperature. For competition ELISAs, blocking IgG was incubated with coated and blocked wells for 1 hour at room temperature. Wells were incubated with peptide-phage in PBS, 0.5% BSA for 1 hour. Plates were washed, incubated with anti-M13-HRP antibody (Sino Biological, catalog number 11973-MM05T-H) and developed with TMB substrate (Mandel, catalog number KP-50-76-03).

### Biolayer interferometry

For IgGs and peptide-IgG fusions, binding kinetics for the S-protein were determined by BLI with an Octet HTX instrument (ForteBio), as described^20^. For peptides, biotinylated peptides (Table 1) were immobilized on streptavidin-coated sensors that were subsequently blocked with excess biotin. Following equilibration with assay buffer, loaded biosensors were dipped for 600 seconds into wells containing 3-fold serial dilutions of S-protein ECD, and subsequently, were transferred back into assay buffer for 600 seconds. Binding response data were corrected by subtraction of response from a reference and were fitted with a 1:1 binding model using the ForteBio Octet Systems software 9.0. We determined K_D_ values at various peptide loading densities to ensure that high densities were not impacting kinetic measurements at the sensor.

### Sequence alignment and analysis

Sequences of binding peptides were imported into Geneious R9 software (Biomatters Ltd.). Peptides were sorted based on the library from which they originated and aligned separately using the MUSCLE algorithm. In the case of peptides lacking cysteines, a strict penalty was imposed on the formation of gaps during alignment. Sequence logos were created using the weblogo server (https://weblogo.berkeley.edu/logo.cgi).

### Virus neutralization assay

The virus neutralization assay was performed as described^20, 24^. Briefly, serial dilutions of IgGs were incubated with the virus at 37 °C for 1 hour. The IgG-virus mixture was transferred to 96-well tissue culture plates containing sub-confluent monolayers of Vero E6 cells in duplicate and incubated at 37 °C and 5% CO_2_. Infection was monitored in one of two ways. The first used a crystal violet based method^24^ as follows. Supernatants were carefully discarded and 100 μl Crystal Violet solution containing 4% formaldehyde was added to each well. Cells were washed and 100 μl containing 50 parts of absolute ethanol (Sigma, St. Louis, MO), 49 parts of MilliQ water and 1 part of glacial acetic acid (Sigma, catalog number 64-19-7) were added to each well. Plates were incubated for 15 minutes at room temperature and read by a spectrophotometer at 590 nm. In addition, we monitored infection using an ELISA-based method^20^ with the anti-S-protein antibody CR3022^25^, followed by incubation with HRP-conjugated goat anti-Fc antibody (Invitrogen, catalog number A18817). Stained cells were visualized using KPL TrueBlue peroxidase (SeraCare, catalog number 5510) and quantified on an ImmunoSpot microanalyzer (Cellular Technologies). All data were analyzed using Prism software (GraphPad Prism 8.0).

### Structural Analysis

The model of two 15033-7 Fabs bound to the SARS-CoV-2 S-protein (PDB entry 7KMK) was imported into PyMol (DeLano Scientific, LLC). Distances between the HC N-termini and the 15033-7 epitope on the unbound RBD were measured as the distance between the C_α_ of the first residue of the HC and the C_β_ of Tyr^489^ of the S-protein. To build the model of two RGN10933 Fabs bound to the S-protein, the model of RGN10933 and RGN10987 bound to the RBD (PDB entry 6XDG) was imported into PyMol along with the data from PDB entry 7KMK. The data from PDB entry 6XDG were duplicated and the RBDs of the model were superposed with the two RBDs in the “up” position in the model from PDB entry 7KMK. The RBDs in the model from PDB entry 6XDG, RGN10987 Fabs, and 15033-7 Fabs were then eliminated from the model, leaving only the two RNG10933 from PDB entry 6XDG bound to the S-protein from PDB entry 7KMK. Distances were measured the same way as for 15033-7.

## Acknowledgements

We are grateful to Daniele Lapa (INMI), for his contribution in determining the neutralizing power of the antibodies tested in Rome. We are also grateful to Carlo Tomino for his valuable help in preparing for the regulatory aspects of <u>monoclonal antibodies</u> in Italy.

## Nomenclature

SARS-CoV-2: severe acute respiratory syndrome coronavirus-2
S-protein: Spike protein
RBD: receptor binding domain
NTD: N-terminal domain
ECD: extracellular domain
VoC: variants of concern
ACE2: angiotensinogen converting enzyme 2
R1-R4: RBD binding peptide 1-4
N1-N4: NTD binding peptide 1-4
nAbs: neutralizing antibodies
HC: IgG heavy chain
LC: IgG light chain
33: neutralizing antibody 15033
33-7: neutralizing antibody 15033-7
NAV: neutravidin

